# Human airway ex vivo models: new tools to study the airway epithelial cell response to SARS-CoV-2 infection

**DOI:** 10.1101/2023.04.15.536998

**Authors:** Said Assou, Engi Ahmed, Lisa Morichon, Amel Nasri, Florent Foisset, Carine Bourdais, Nathalie Gros, Sonia Wong, Aurelie Petit, Isabelle Vachier, Delphine Muriaux, Arnaud Bourdin, John De Vos

## Abstract

Airway-liquid interface cultures of primary epithelial cells and of induced pluripotent stem cell-derived airway epithelial cells (ALI and iALI, respectively) are physiologically relevant models for respiratory virus infection studies because they can mimic the *in vivo* human bronchial epithelium. Here, we investigated gene expression profiles in human airway cultures (ALI and iALI models) infected or not with severe acute respiratory syndrome coronavirus 2 (SARS-CoV-2) using publicly available and our own bulk and single-cell transcriptome datasets. SARS-CoV-2 infection significantly increased the expression of interferon-stimulated genes (*IFI44*, *IFIT1*, *IFIT3*, *IFI35*, *IRF9*, *MX1*, *OAS1*, *OAS3* and *ISG15*) and inflammatory genes (*NFKBIA*, *CSF1*, *FOSL1*, *IL32* and *CXCL10*) at day 4 post-infection, indicating activation of the interferon and immune responses to the virus. Extracellular matrix genes (*ITGB6*, *ITGB1* and *GJA1*) also were altered in infected cells. Single-cell RNA sequencing data revealed that SARS-CoV-2 infection damaged the respiratory epithelium, particularly mature ciliated cells. The expression of genes encoding intercellular communication and adhesion proteins also was deregulated, suggesting a mechanism to promote shedding of infected epithelial cells. These data demonstrate that ALI/iALI models help to understand the airway epithelium response to SARS-CoV-2 infection and are a key tool for developing COVID-19 treatments.

## Introduction

The rapid spread of Severe Acute Respiratory Syndrome Coronavirus 2 (SARS-CoV-2) in humans has posed a serious global health threat. Coronaviruses are part of a large family of viruses that cause illnesses ranging from common colds to severe respiratory diseases, including COVID-19 caused by SARS-CoV-2. SARS-CoV-2 is a single positive-stranded RNA enveloped virus that replicates in epithelial cells [1]. Upon infection, SARS-CoV-2 binds the host ACE2 receptor through its spike protein, and enters the cells by fusion of the viral membrane with the epithelial cell membrane or by endocytosis [2]. After binding, the spike protein can be cleaved by TMPRSS2, a host membrane serine protease, facilitating viral entry [1]. Then, the virus replicates inside epithelial cells and produces newly synthetized viral particles that are secreted by the host cells [3].

Due to the COVID-19 long- and short-term effects on human health and the need to limit the emergence of novel virus variants, significant efforts have been dedicated to understand the viral infection mechanisms and to develop antiviral drugs using physiologically relevant *in vitro* culture models that mimic *in vivo* phenotypes. Human airway epithelial cells in culture are traditionally used for modeling respiratory diseases [4]. These cells can be obtained from lung tissue biopsies and are cultured as primary airway epithelial cells in air-liquid interface (ALI) systems that support epithelial cell differentiation and mimic key aspects of the mucosal epithelium [5]. They can also be derived by differentiation of induced pluripotent stem cells (iPSCs) in ALI systems (i.e. iALI) [6–9]. In both ALI and iALI systems, epithelial cells are cultured on a permeable membrane with the medium in the basal chamber and the epithelium exposed to air at the apical side of the membrane. In this system, cells are in contact with air and can be induced to differentiate into a functional pseudo-stratified epithelium. Several studies confirmed that ALI culture transcriptomic profiles are very similar to those of *in vivo* airway epithelium obtained by bronchial brushing or biopsy [10,11]. ALI and iALI models can be infected by viruses and have been used to model various mechanisms of viral pathogenesis [12,13]. SARS-CoV-2 can replicate in both models [1,14] and infection can be limited by interferon. This demonstrated interferon therapeutic potential for COVID-19 treatments and the usefulness of these models as a high-throughput screening tool [15,16]. Additionally, ALI and iALI cultures from patients with respiratory diseases (e.g. chronic obstructive pulmonary disease) recapitulate some of the *in vivo* disease characteristics, and are used to assess the impact of smoke exposure on viral infection [12,17,18]. Therefore, ALI and iALI cultures could help to understand SARS-CoV-2 effects in bronchial epithelium, by analyzing the transcriptomic changes upon infection. ALI models are very helpful to identify the key initiating steps of viral injury and innate epithelial cell defense that may or may not lead to cell infection and replication. Increasing the number of models and of donors will improve the robustness of the identified pathways by reducing the inter-individual heterogeneity in viral susceptibility.

Various omic-based studies, including *in vivo* (human samples) and *in vitro* (model systems) transcriptome profiling studies (bulk RNA sequencing), have highlighted the molecular changes induced by SARS-CoV-2 infection [19–21]. The recent advent of single-cell RNA-sequencing (scRNA-seq) provides a precious approach to carefully analyze gene expression and cell composition. For instance, scRNA-seq has been used to identify the cells in the human respiratory system with the highest expression of transmembrane receptors for SARS-CoV-2 [22], and to show that in ALI cultures of nasal epithelial cells, ciliated and goblet/secretory cells express progressively SARS-CoV-2 entry factors. Moreover, in infected human bronchial epithelial–derived ALI cultures, scRNA-seq [23] revealed that ciliated cells are a major SARS-CoV-2 target.

In the present study, we analyzed bulk and single-cell RNA-seq datasets to provide a detailed picture of the gene expression changes in ALI and iALI models following SARS-CoV-2 infection. This analysis highlighted the molecular mechanisms involved in the induction of the hyper-inflammatory state and innate immune response, including interferon signaling, chemokines and extracellular matrix (ECM), and identified potential regulatory microRNAs (miRNAs) for therapeutic interventions.

## Methods

### ALI culture of primary airway epithelial cells and iPSC-derived airway epithelium for scRNA-seq analysis

Primary Human Bronchial Epithelial Cells (HBEC) were expanded and differentiated in ALI culture following the protocol given by StemCell Technologies. Briefly, cells were dissociated mechanically from bronchial biopsy specimens obtained by fiberoptic bronchoscopy (approval number: 2013 11 05; NCT02354677) and cultured in PneumaCult-Ex Plus expansion medium (cat #05041, StemCell Technologies, France) for 15 days. After the expansion phase, differentiation was initiated by seeding 1.1 × 10^5^ cells/insert on Transwell^R^ polyester membranes (ref 3460, Corning, Kennebunk, United States). Once epithelial cells reached confluence, the apical growth medium was removed, and the basal medium was replaced by PneumaCult™-ALI maintenance medium (cat #05002, StemCell Technologies, France) (i.e. day 0 of ALI culture). Epithelial cells were allowed to differentiate at 37°C, 5% CO_2_ for 28 days.

iPSC-derived airway epithelium on ALI (iALI model) was generated as previously described [6]. Briefly, the major stages of embryonic lung development were recapitulated as follows: stage 1, definitive endoderm (day 0–3) using activin A; stage 2, anterior foregut endoderm (day 4–8); stage 3, lung progenitor specification (day 9); and stage 4, epithelial layer (day 14). After 40 days, the airway epithelium on iALI displayed morphologic and functional similarities with primary human airway epithelial cells and included different airway cell types (basal, secretory, and multi-ciliated cells).

### ScRNA-seq and data analysis

Non-infected ALI and iALI cultures were dissociated with trypsin into single-cell suspensions. Cell viability and aggregation were tested before starting the single-cell library preparation. The concentration of freshly dissociated cells was adjusted to 1000□cells/μl in HBSS/0.05% BSA and then the 10x Chromium Controller and the Chromium Single Cell 3’ Reagent kit V3.1 were used for performing the scRNAseq experiments. Library preparation was performed according to the manufacturer’s instructions using the Chromium Chip B Single Cell kit, and Chromium Multiplex Kit (10X Genomics). Sequencing was performed in paired-end mode with an S1 flow cell (28/8/87 cycles) and a NovaSeq 6000 sequencer (Illumina) at the MGX core facility of Montpellier, France. First, the cell ranger mkfastq and cellranger count pipelines were used for the initial quality control, sample demultiplexing, mapping, and raw data quantification. Briefly, fastq files were run with the Count application using default parameters and were aligned to the human genome reference sequence GRCh38, filtered and counted. The C-loop software (version 6.2.0) was used to visualize clusters and sub-clusters of transcriptionally related cells and to identify candidate genes the expression of which was enriched in specific clusters. Clustering results were visualized with the t-distributed Stochastic Neighbor Embedding (tSNE) technique. Bronchial cell (biopsy/brushing samples) datasets included in [24] were analyzed through a COVID-19 Cell Atlas website portal and were visualized using Uniform Manifold Approximation and Projection (UMAP).

### Functional enrichment analysis of bulk RNA-seq datasets

The unique bulk RNA-seq signature was obtained from a publicly available list of differential expressed genes between primary human lung epithelium infected with SARS-CoV-2 (multiplicity of infection, MOI, = 2 for 24 h) and mock-treated controls [20]. The GO functional enrichment and pathway enrichment analyses were performed with ShinyGO [26] and Gene Set Enrichment Analysis (GSEA) (http://www.broadinstitute.org/gsea/). GO annotations were divided in three categories: biological process, molecular function, and cellular component. In the enrichment analysis, the Fisher’ exact test was used to test whether genes were enriched in a term, and an adjusted *p* value <0.05 was set as the screening condition. The GenGo Metacore software was used to identify miRNA targets. Heatmaps and gene networks were generated with Ingenuity Pathway Analysis (IPA, QIAGEN, Redwood City, CA, USA).

### Analysis of publicly available single-cell RNA-seq datasets

A public scRNA-seq dataset (GEO accession number: GSE166766) of HBEC ALI culture samples infected with SARS-CoV-2 (MOI ∼0.01) was also analyzed [25]. The Velocyto® package was used to investigate the gene expression dynamics in the scRNA-seq data before virus infection and at day 3 post-infection (dpi). RNA velocity quantifies the change in the state of a cell over time by distinguishing unspliced and spliced mRNAs reads. To obtain de counts of spliced and unspliced mRNAs reads, Velocyto used the outputs of CellRanger (version 3.0.2) alignments with the command line ‘run10x’ and the transcriptome GRCh38.p12 (accession NCBI:GCA_000001405.27). At this step, loom files were created for the input data of the scVelo tool [26]. This python package allowed normalizing with the scv.pp.normalize_per_cell() function and transform scv.pp.log1p(). Genes were filtered by keeping only the 2000 top highly variable genes (n_top_genes = 2000 parameters for scv.pp.filter_and_normalize function). Then, moments were calculated for each cell across its nearest neighbors with n_neighbors set to 30 and the first 10 PCs using the scv.pp.moments() function. Then, cell velocities were estimated using scvelo.tl.velocity() based on a stochastic model of transcriptional dynamics. To visualize the velocity graph, data were projected on the UMAP space and clusters defined by the CellRanger pipeline were colored. After cluster identification on Seurat V4 [27], the function cluster_analysis from the SingleCellSignalR package [28] was used to compute paracrine interactions between cell clusters and to predict ligand–target links between interacting cells by combining their expression level with prior knowledge on gene regulatory networks and signaling pathways. To compare ALI and iALI samples, genes identified in our samples were combined with other publicly available datasets that used infected iPSC-derived AT2 cells (iAT2) [16], specifically for the genes related to inflammatory and interferon responses and extracellular ECM.

### SARS-CoV-2 virus stock and titration

The hCoV-19/France/HDF-IPP11602i/2021 (21A - Delta – B.1.617.2) strain was supplied by the National Reference Centre for Respiratory Viruses hosted by Institut Pasteur (Paris, France). The human sample from which this strain was isolated was provided by Dr Guiheneuf Raphaël, CH Simone Veil, Beauvais France. The strain was propagated in VeroE6 cells with DMEM containing 25mM HEPES at 37°C and 5% CO_2_ and viruses were harvested 72 hours post-inoculation. Virus stocks were stored at −80°C. Viruses from infected cell culture supernatants were titrated with the plaque assays on a monolayer of VeroE6 cells and 100µL of virus serial dilutions. The plaque forming unit (PFU) values were determined by crystal violet staining and then by scoring the wells displaying cytopathic effects. The virus titer was determined as the number of PFU/mL, and MOI was the PFU/cell ratio.

### Reverse transcription-quantitative polymerase chain reaction (RT-qPCR)

RNA was extracted from cells using the QIAshredder kit (QIAGEN, Redwood city, CA, USA) and the RNeasy mini kit (Qiagen, Redwood city, CA, USA) according to the manufacturer’s instructions. Viral RNA was quantified by RT-qPCR in triplicate as described [29], using the Luna Universal One-Step RT-qPCR Kit (New England Biolabs, Ipswich, MA, USA) and a BIORAD CFX Opus 384 system. Relative gene expression was calculated for each triplicate by normalizing to *GAPDH* level (control) and using the formula 2^−ΔCt^. Primers are listed in Supplementary Table S1.

### Immunofluorescence analysis

iALI cultures were fixed in 4% paraformaldehyde for 4 hours. After three PBS washes, iALI samples were stored in PBS at 4°C. Samples were permeabilized in 0.5% Triton X-100/PBS at room temperature for 20 min and then blocked with PBS/0.1% Triton X-100/1% bovine serum albumin (BSA)/10% donkey serum at room temperature for at least 1h. Primary antibodies against p63 (AF1916, Biotechne, Minneapolis, MN, USA), TubIV (T7941, Sigma, Saint Louis, MO, USA) and SARS-CoV-2 M protein membrane (100-401-A55, Rockland Immunochemicals, Pottstown, PA, USA) were diluted (1/100, 1/200 and 1/200, respectively) in PBS/1%BSA/0.1%, Triton X-100 and added to the samples for overnight incubation. Then, samples were washed three times with PBS/0.025% Triton X-100 before incubation (room temperature for 2h) with the following secondary antibodies: anti-mouse coupled to Alexafluor 555 (A31570, Invitrogen, Waltham, MA, USA), anti-rabbit coupled to Alexafluor 488 (A21206, Invitrogen, Waltham, MA, USA) and anti-goat coupled to Alexafluor 647 (A21447, Invitrogen, Waltham, MA, USA) (all diluted to 1/1000 in PBS/1% BSA/0.1%, Triton X-100. After three washes in PBS/0.025% Triton X-100, samples were incubated with DAPI (D9542, Sigma, Saint Louis, MO, USA), diluted 1/2500 in PBS, for 5 min and rinsed in PBS. Then, iALI samples were separated for their support and mounted between glass slides. Confocal images were acquired using a Cell-Discoverer 7 LSM900 confocal laser-scanning microscope (Zeiss, Germany) at 40x magnification, and processed with Zen Blue.

### Statistical analysis

Data are presented as the mean ± SEM, unless otherwise specified. Statistical analyses were performed with the GraphPad Prism 5 software (Student’s *t*-test; GraphPad). The shown data were from representative experiments, with similar results in at least three independent biological replicates, unless otherwise specified. Differences were evaluated using the Student’s *t*-test. A *p* value ≤0.05 was considered significant.

## Results

### Cellular landscapes of non-infected human lung epithelium models

One of the challenges in COVID-19 research is the need of physiologically relevant *in vitro* cell culture models to mimic the native environment. To this aim, non-infected HBEC in ALI culture and iPSC-derived epithelial-like cells in iALI culture were analyzed by scRNA-seq to generate a comprehensive repertoire of the cell populations present in these models. Single-cell transcriptome data were obtained also from lung epithelium collected by bronchial biopsy or brushing (Figure 1A). Analysis of these scRNA-seq data from these three models (biopsy/brushing bronchial cells, ALI, and iALI cultures) (Figure 1B) showed that most epithelial cell markers, such as *SCGB1A1/CC10* (secretoglobin family 1A member 1; secretory cells), *TP63* (tumor protein p63; basal cells), and *FOXJ1* (forkhead box J1; multi-ciliated cells) were present in the three sample types. This suggested that non-infected ALI and iALI models contain the essential cell types necessary for bronchial epithelium function and can be used as experimental *ex vivo* models to study SARS-CoV-2 infectivity. Comparison of the single-cell landscapes indicated that unlike the biopsy/brushing and ALI models, the iALI model expressed *ASCL1*, a marker of neuroendocrine cells, but not *FOXI1* (ionocyte marker) and all models express *POU2F3* (tuft cell marker). In addition, immune cells were an important cell fraction in biopsy/brushing samples (Figure 1C), but were absent in the ALI and iALI models. This major difference may help to (i) understand the epithelial-intrinsic innate inflammatory response to SARS-CoV-2 infection; (ii) identify the key epithelial target cells; and (iii) compare the molecular mechanisms triggered in the absence (iALI and ALI) and the presence (biopsy–brushing samples) of immune cells.

**Figure 1:**
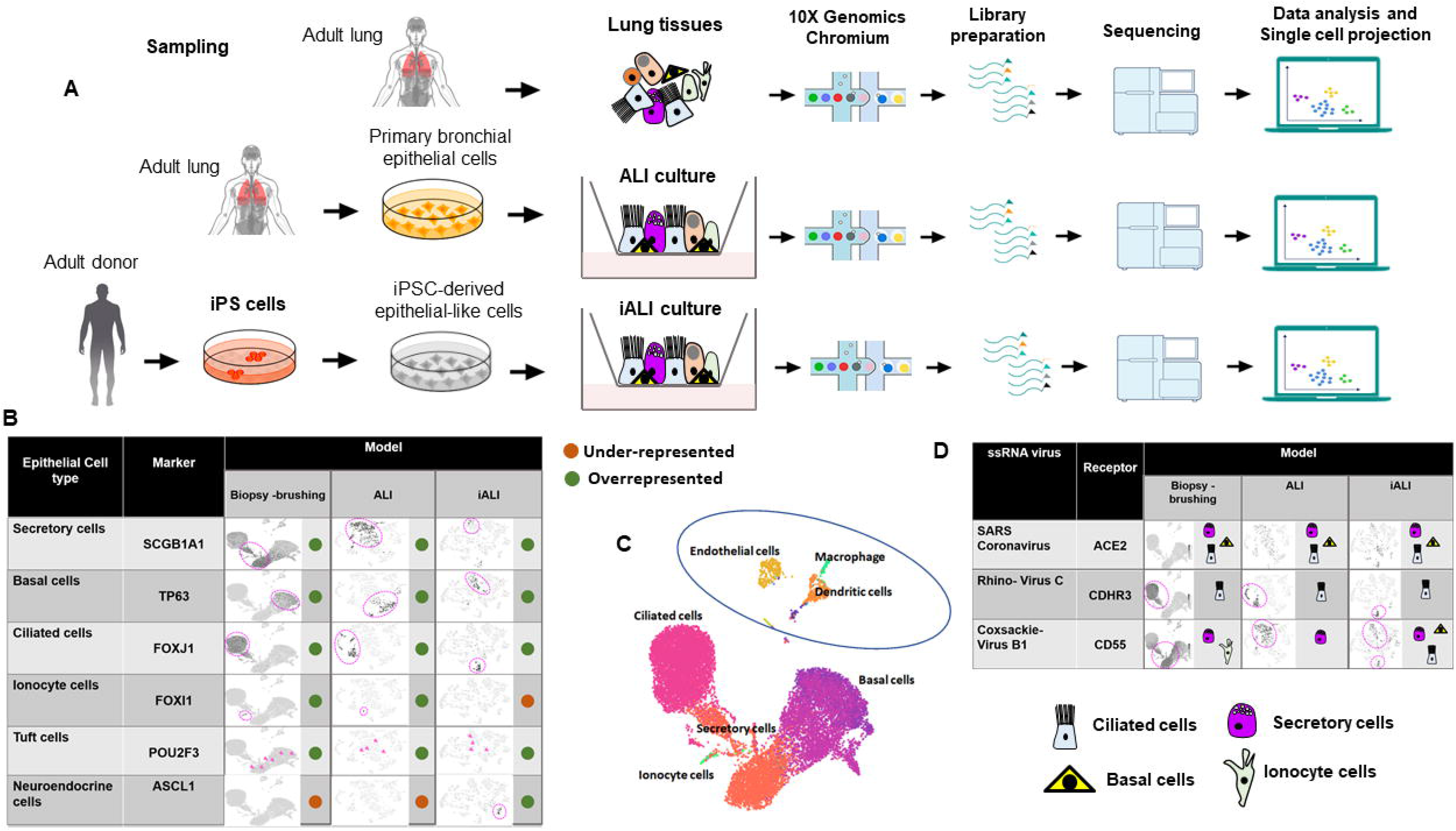
Distribution of the different airway epithelial cell types the three lung tissue models. (A) Schematic representation of the scRNA-seq experimental workflow: airway epithelium sources (biopsy/brushing-derived cells, ALI culture of primary bronchial epithelial cells and iALI culture of iPSC-derived epithelial-like cells), generation of scRNA-seq libraries and sequencing, computational analysis to identify cell types. The brushing/biopsy epithelium scRNA-seq data were from [24]. The ALI and iALI scRNA-seq data were generated in our laboratory (see Methods). (B) Contribution of each cell type in the three models. UMAP and tSNE were used to show the contribution of *SCGB1A1*+, *TP63*+, *FOXJ1*+, *FOXI1*, *POU2F3*+, and *ASCL1*+ cells (dark gray). (C) UMAP projection of the different cell types. The feature plots display the subset of airway epithelial cells obtained from biopsy/brushing of healthy donors [24]. (D) Maps of the expression of ssRNA virus receptors in single cells from the three models. UMAP and tSNE were used to show the cell types that express the ACE2, CDHR3 and CD55 receptors.

### Cell-specific expression of single-stranded RNA virus receptors in non-infected human lung epithelium models

The expression of receptors to which single-stranded RNA (ssRNA) viruses can bind is a key factor for virus infection and transmission. Analysis of scRNA-seq data from the three non-infected lung epithelium models showed that *CDHR3* (encoding cadherin related family member 3, the entry receptor for rhino-virus C) [30] was specifically expressed in ciliated cells, and *CD55* (Coxsackie-virus B1 receptor) [30] in secretory cells (Figure 1D). *ACE2* (SARS-CoV-2 receptor) was expressed by ciliated, secretory and basal cells. Moreover, in biopsy/brushing samples, ACE2^+^ cells were overrepresented among all epithelial cell types compared with immune cells, including macrophage, endothelial and dendritic cells (Figure 1D). This is in line with a recent study reporting the absence of viral transcripts in bronchoalveolar fluid and peripheral blood mononuclear cell samples collected from patients with COVID-19 [31]. Collectively, scRNA-seq data analysis of the three cell culture models showed expression of ssRNA virus receptors in secretory, ciliated and basal cells, suggesting the possibility of modeling ssRNA virus infection, including SARS-CoV-2, in airway epithelium *in vitro*.

### Changes induced by SARS-CoV-2 infection in the ALI model

A recent bulk RNA-seq dataset [20] was used to compare the gene expression profiles of infected and non-infected human bronchial epithelium in order to identify the “SARS-CoV-2 bulk RNA-seq signature” induced by SARS-CoV-2 infection in the ALI model (Supplementary Table S2). GO biological process and molecular function analyses of this “SARS-CoV-2 bulk RNA-seq signature” showed significant overexpression of genes implicated in immune response, cytokines and chemokine activity (Figure 2A). The GO cellular component enrichment analysis showed that many differentially expressed genes were related to ECM and ECM regulators (Figure 2A). GSEA confirmed the significant upregulation of the interferon and inflammatory pathways (Figure 2B). Heatmaps revealed that the core transcriptional response included genes implicated in “immune cell trafficking”, “inflammatory response”, “cellular movement”, “inflammatory response” and “cell-to-cell signaling and interaction” (Figure 2C), which is consistent with the biological processes and molecular functions highlighted above. The top categories, ranked in accordance with their −log(P-value), are shown in Supplementary Table S3. In total, 146 enriched canonical pathways were identified (Supplementary Table S4). The interferon signaling pathway was ranked first with a −log (P-value) of 14.1. Many genes belonging to the interferon machinery implicated in the antiviral response and viral replication were identified (Figure 2D), such as interferon-induced genes (*IFI27*, *IFI35*, *IFI44*, *IFI44L*, *IFIH1*, *IFIT1*, *IFIT3*, *IFITM1*, *IFITM3*), interferon regulatory factors (*IRF7* and *IRF9*), interferon-stimulated genes (*ISG15* and *ISG20*), MX dynamin-like GTPases (*MX1* and *MX2*), oligoadenylate synthetases (*OAS1*, *OAS2* and *OAS3*) and the master suppressor of cytokine signaling (*SOCS3*). With a Z-score >2 as a threshold of significant activation, the following signaling pathways were identified as activated: ‘*IL17* signaling’ (Z-score=5.099), ‘*HMGB1* signaling’ (Z-score=4), ‘*TREM1* signaling’ (Z-score=3.742), ‘Toll-like Receptor signaling’ (Z-score=2.53), ‘*IL-8* signaling’ (Z-score=4.123), ‘iNOS signaling’ (Z-score=3.162), ‘*IL-6* signaling’ (Z-score=3.37), ‘*TNFR1* signaling’ (Z-score=2.121), ‘*IL-1* signaling’ (Z-score=3) and ‘*PI3K/AKT* signaling’ (Z-score=2.333). Conversely, some of the enriched signaling pathways had Z-scores lower than - 2, such as ‘inhibition of matrix metalloproteases’ (Z-score=-2), ‘*PPAR* signaling’ (Z-score=-3.606) and ‘erythropoietin signaling pathway’ (Z-score=-2.524). Concerning upstream regulators (Supplementary Figure S1, Supplementary Table S5), the highest ranked regulatory effects with consistency scores up to 4.8 strongly suggested that the upstream regulators *TNF*, *IL1A* and *F2* may be responsible for the gene expression changes in the SARS-CoV-2 bulk RNA-seq signature. IPA analysis predicted that these upstream regulators are involved in chemotaxis, invasion and cell movement, mainly through induction of their targets, including many genes associated with the innate immune response (*IL1B*, *NFKBIA*, *CXCL1*, *NOD2*, *PLAUR*, *C3*, *ITGB2*, *S100A8*, *PLAU*, *S100A9*, *IL1A*, *TNF*), cytokine and chemokine activities (*CCL20*, *CXCL2*, *CXCL3*, *CXCL5*, *CXCL6*, *CXCL8*, *CXCL10*, *IL6*, *IL8*, *IL32*, *IL33*, *CSF3*, *CSF2*, *CSF1*), and genes implicated in ECM degradation, such as matrix metalloproteases (*MMP13*, *MMP9*, *MMP1*). These metalloproteases are connected to pro-inflammatory chemokines and play an operative role in ECM degradation during inflammation that can be triggered by virus infection. Our data highlighted and confirmed that matrix metalloproteinases play key roles in viral infection and its progression [32] through airway remodeling (i.e. loss of the epithelium barrier integrity [33] and elastin degradation in the ECM [34]).

**Figure 2:**
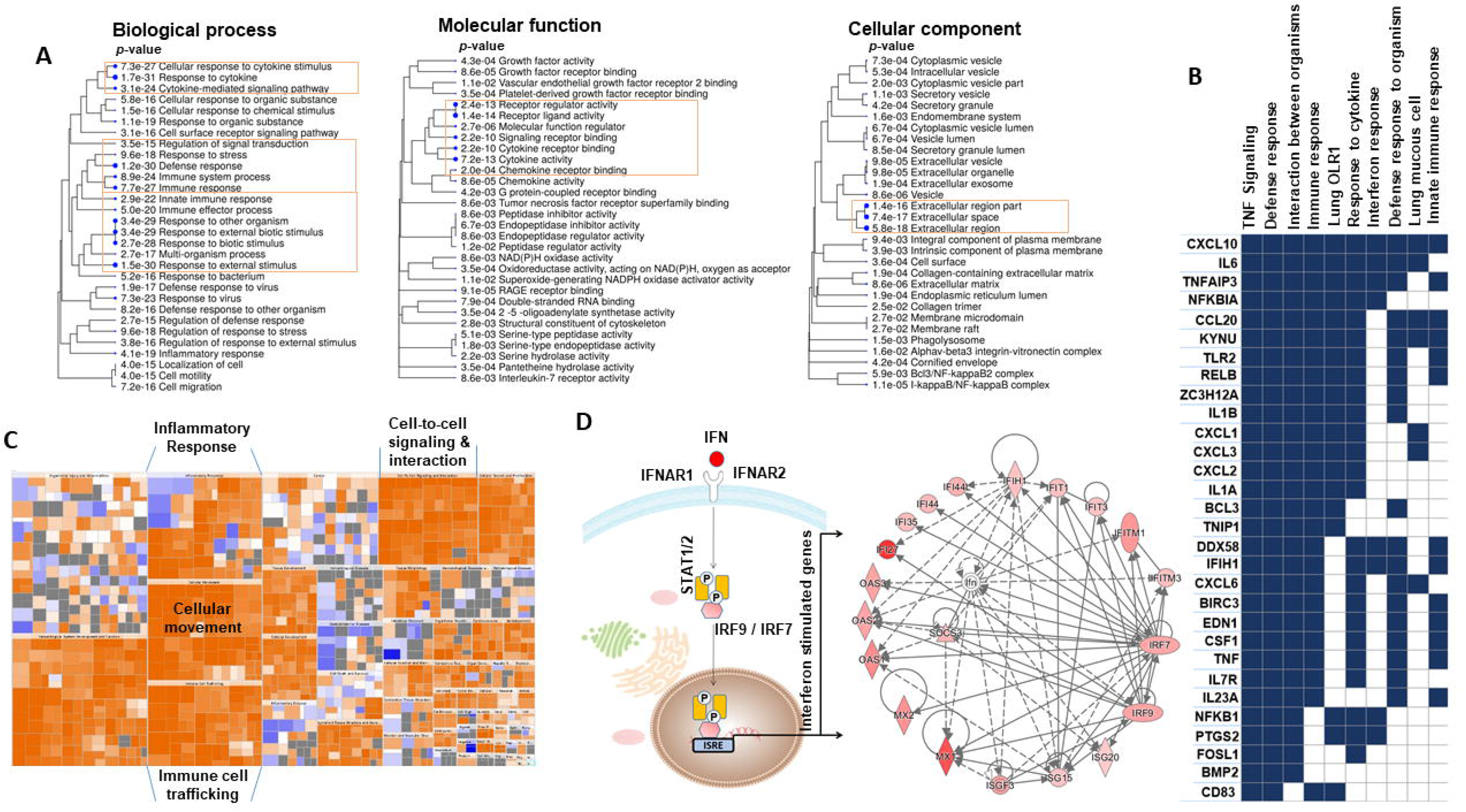
Analysis of GO terms enriched in the unique bulk RNA-seq signature of the ALI model upon SARS-CoV-2 infection. (A) Pathway enrichment analyses were performed with human gene names. The size of the blue dots corresponds to the enrichment (FDR) and bigger dots indicate more significant *p*-values. The biological process, molecular function, and cellular component categories revealed the high enrichment of immune response and cytokine/chemokine activities upon SARS-CoV-2 infection. (B) GSEA performed using the unique bulk RNA-seq signature upon SARS-CoV-2 infection. The heat map shows the (clustered) genes in the leading-edge subsets and the dynamic expression of genes involved in immune response, interferon response, defense response, TNF-signaling and response to cytokines. (C) Enrichment heat map (IPA) showing the dynamic activity of canonical pathways after SARS-CoV-2 infection. Each colored rectangle is a biological function and the color range indicates its activation state (orange for an activated pathway with Z-score > 2, and blue for an inhibited pathway with Z-score <−2). The pathways were classified into different types according to the IPA database. (D) The network shows the interactions of interferon (IFN)-stimulated genes. Nodes shaded in pink represent protein-coding genes that are upregulated in the ALI model upon SARS-CoV-2 infection. Labels in nodes and edges (lines) illustrate the nature of the relationship between genes and their functions. A dotted line represents an indirect interaction and a solid line a direct interaction. IPA, Ingenuity Pathway Analysis. GO, Gene Ontology.

### Conserved expression of the epithelial cell response to SARS-CoV-2 infection in the iALI model

Then, bulk RNA sequencing datasets of iPSC-derived alveolar epithelial type 2-like cells (iAT2) infected by SARS-CoV-2 and mock-infected [16] at 1 and 4 dpi were analyzed to determine whether the SARS-CoV-2 bulk RNA-seq signature observed in the ALI model was present also in the iALI model. Analysis of the temporal distribution of RNA-seq reads allowed identifying four major gene groups (Figure 3A): (a) genes that were upregulated early (at 1 dpi) and the expression of which gradually increased from 1 to 4 dpi (*NFKBIA*, *CSF1*, *IL32* and *FOSL1)*; (b) genes that were upregulated early and the expression of which was not changed at 4 dpi (*IL23A*, *CXCL10*, *CXCL20* and *PLAUR)*; (c) genes that were upregulated at 1 dpi and were then downregulated at 4 dpi (*CXCL3*, *CXCL5*, *CXCL1* and *LOX)*; and (d) genes that were upregulated only at 4 dpi (*MMP9* and *MMP13*). In addition, and like in the ALI model, in the COVID-19 iALI model, the interferon response was activated, as indicated by the upregulation of genes encoding interferon induced proteins and interferon regulatory factors (Figure 3B). This suggests that the ALI and iALI models share common interferon response regulation features. Conversely, the expression kinetics of ECM-related genes showed that some genes (*ITGB6*, *ITGB1*, *GJA1*, *VIM* and the ECM regulator *PLOD2*, which encode proteins of the host cell cytoskeleton structure) were progressively downregulated during SARS-CoV-2 infection in the iALI model (Figure 3C). This indicated that the ECM and adhesion pathways are affected during cell-to-cell SARS-CoV-2 transmission. Moreover, the expression of *EpCAM* (epithelial cell adhesion molecule), an epithelial cell surface marker, progressively decreased, and that of the stromal cell marker *COL1A1* progressively increased (Figure 3D), suggesting epithelial mesenchymal transition.

**Figure 3:**
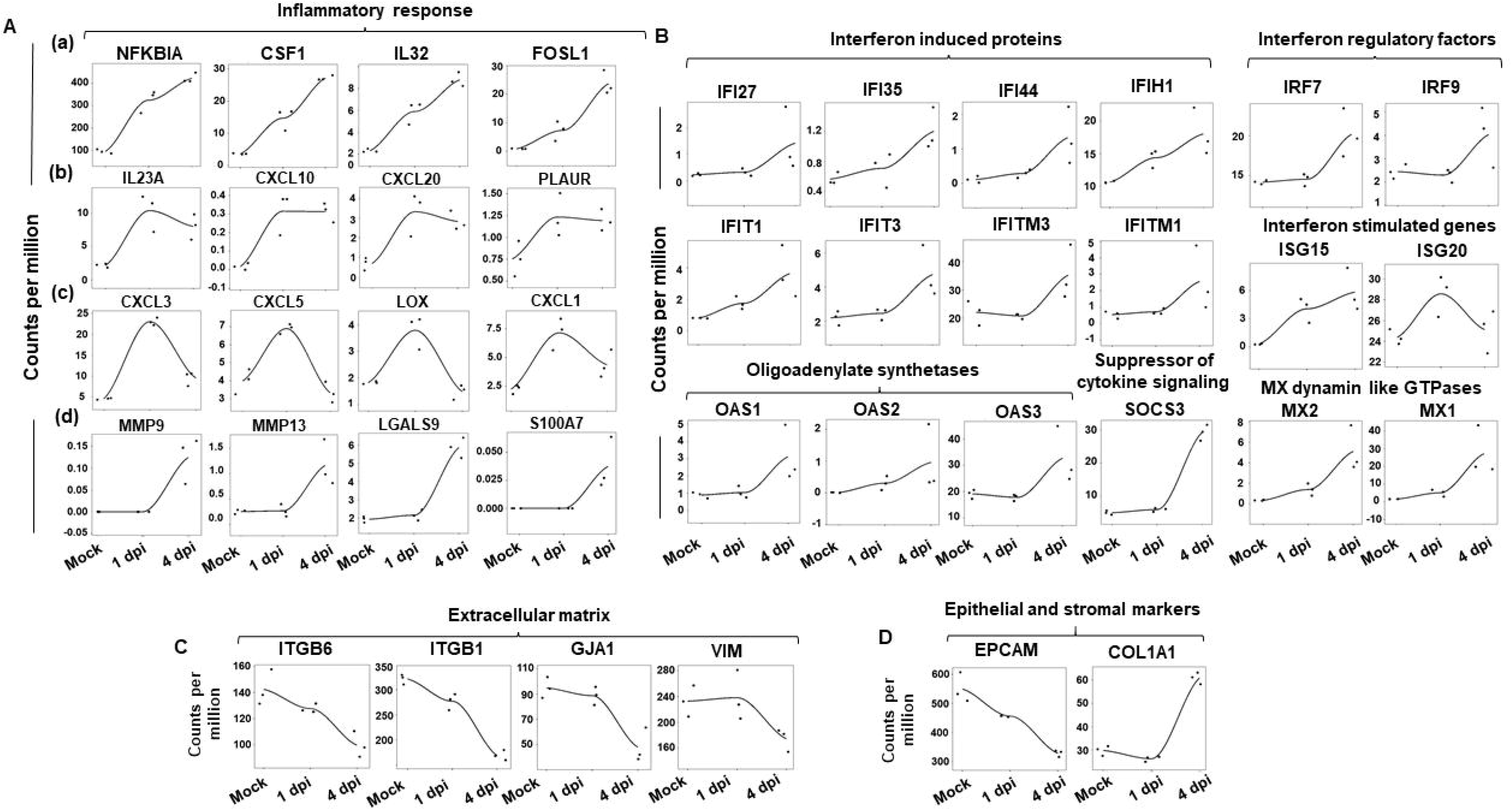
Dynamic gene expression changes in the iALI model upon SARS-CoV-2 infection. Expression levels of (A) inflammatory cytokines/chemokines, including (a) genes that were upregulated early and the expression of which gradually increased upon SARS-CoV-2 infection, from 1 to 4 dpi; (b) genes that were upregulated early and the expression of which remained constant between 1 and 4 dpi; (c) genes that were upregulated specifically at 1 dpi and were then downregulated at 4 dpi; and (d) genes that became upregulated at 4 dpi. Expression of genes implicated in the (B) interferon response, (C) extracellular matrix, and (D) epithelial/stromal gene upon. Normalized expression levels (counts per million reads) were quantified by RNA-seq using data from [16] (human iPSC-derived AT2 cells infected or not with SARS-CoV-2).

### Experimental validation of the epithelial cell response to SARS-CoV-2 infection in the iALI model

To gain insight into the molecular basis of SARS-CoV-2 infection in the iALI model, iALI bronchial epithelium was exposed to SARS-CoV-2 Delta at low MOI (0.05) to let the infection take hold for 4 days and then the expression of interferon-induced genes and regulatory factors, viral entry genes and ECM genes was investigated. Viral DNA quantification confirmed infection of iALI cultures. The virus was detected at the apical side at 1 dpi and also at 4 dpi (Figure 4A). Moreover, SARS-CoV-2 membrane protein colocalized with tubulin α 4a (TubIV), suggesting preferential infection of ciliated cells (Figure 4B). RT-qPCR analysis of infected cells showed activation of the interferon signaling pathways at 1 dpi and 4 dpi. Interferon-induced proteins (*IFI44*, *IFIT1*, *IFIT3*, *IFI35*), interferon regulatory factors (*IRF9*, *MX1*, *ISG15*), oligoadenylate synthetases (*OAS1*, *OAS3*) and chemokine ligand (*CXCL10*) were upregulated at 4 dpi (Figure 4C). Conversely, viral infection did not alter the expression of genes encoding adhesion molecules, such as *GJA1* and *ITGB1*, unlike what observed upon infection at higher MOI (0.5) (Figure 3C). In agreement with this observation, trans-barrier electrical resistance (TEER) of iPSC-derived epithelial cells was comparable in non-infected and infected (at low MOI = 0.05) iALI cultures from 1 to 4 dpi, indicating that at this infection level, epithelium integrity was not disrupted (Figure 4D). Altogether, these data show that the iALI model is a faithful and sensitive model for SARS-CoV-2 infection.

**Figure 4:**
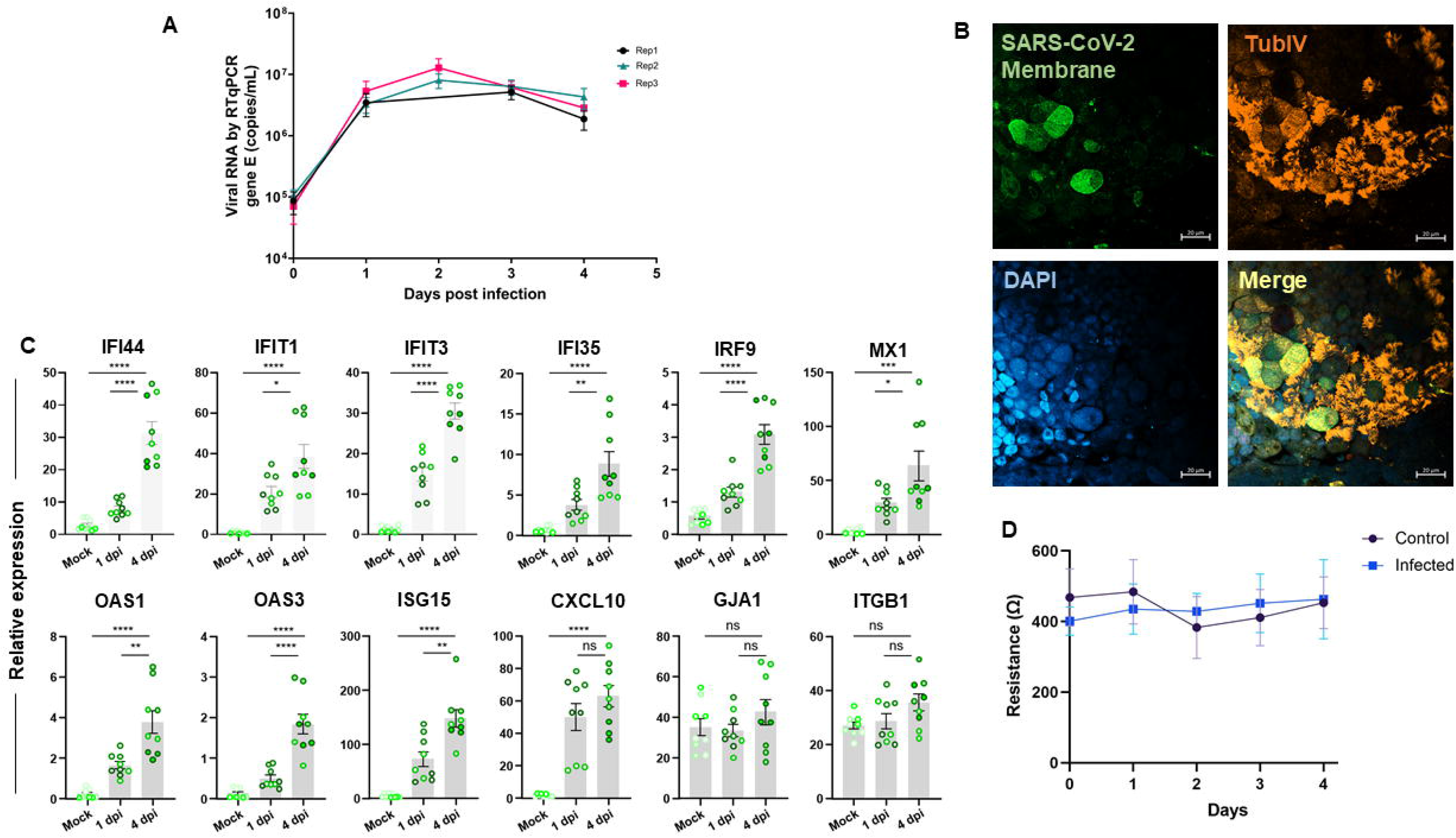
Validation of SARS-CoV-2 infection in the iALI model. (A) iALI cultures were infected with SARS-CoV-2 Delta (MOI=0.05) and viral RNA was quantified by RT-qPCR (gene E/mL). iALI cultures were washed by adding culture medium at the apical side at 37°C for 15 min, then viral RNA was extracted and quantified in triplicate by RT-qPCR (amplification of the viral envelope gene). Data are from three independent experiment (a, b, c) using three different iALI cultures. The standard deviation shows the result variability. (B) iALI cultures were stained with anti-SARS-CoV-2 M protein (viral membrane, green), anti-α-tubulin (ciliated cell marker, orange) and anti-P63 (basal cell marker, red) antibodies. Nuclei were counterstained with DAPI (blue). SARS-CoV-2 was identified on the motile cilia. Scale bar: 20 µm. (C) RT-qPCR analysis of the expression of genes encoding inflammatory and interferon-related factors in iALI cultures infected with SARS-CoV-2 (MOI 0.05) at 1 dpi and 4 dpi. Data are the mean□±□SEM of three independent experiments with three technical replicates/each (9 samples); ^∗^p <0.05, ∗∗p <0.01, ∗∗∗p <0.001, ∗∗∗∗p <0.0001, ns: not significant (Student’s *t* test). (D) Transepithelial resistance was measured daily with an EVOM2 (WPI, Friedberg, Germany), while the apical side was submerged with culture medium. Measurement were done after 10 min of incubation at 37°C. MOI: multiplicity of cellular infection.

### Epithelial cell communication networks in response to SARS-CoV2 infection

ALI cultures include different cell types connected by tight and adherens junctions. Communication between epithelial cells occurs through the release of a variety of small molecules, including cytokines and chemokines. To investigate the impact of SARS-CoV2 infection on ligand/receptor interaction between the different airway cell types, scRNA-seq data from HBEC ALI cultures infected or not with SARS-CoV-2 (MOI 0.01) were analyzed using the SingleCellSignalR method. Comparison of the summary chord diagram indicated a decrease in the number of paracrine interactions between the main epithelial cell types after 2dpi (Figure 5A), which strongly suggests an impact of SARS-CoV2 infection on intercellular communications. For instance, expression of receptor tyrosine kinase (*RET*) and its ligand artemin (*ARTN*), one of the most preeminent interacting pair between neuroendocrine cells and other epithelial cells, was strongly decreased at 3 dpi (Figure 5B). This suggests that neuroendocrine cells are particularly sensitive to environmental stimuli (e.g., viral infection), and act as a rheostat to orchestrate ALI-culture responses.

**Figure 5:**
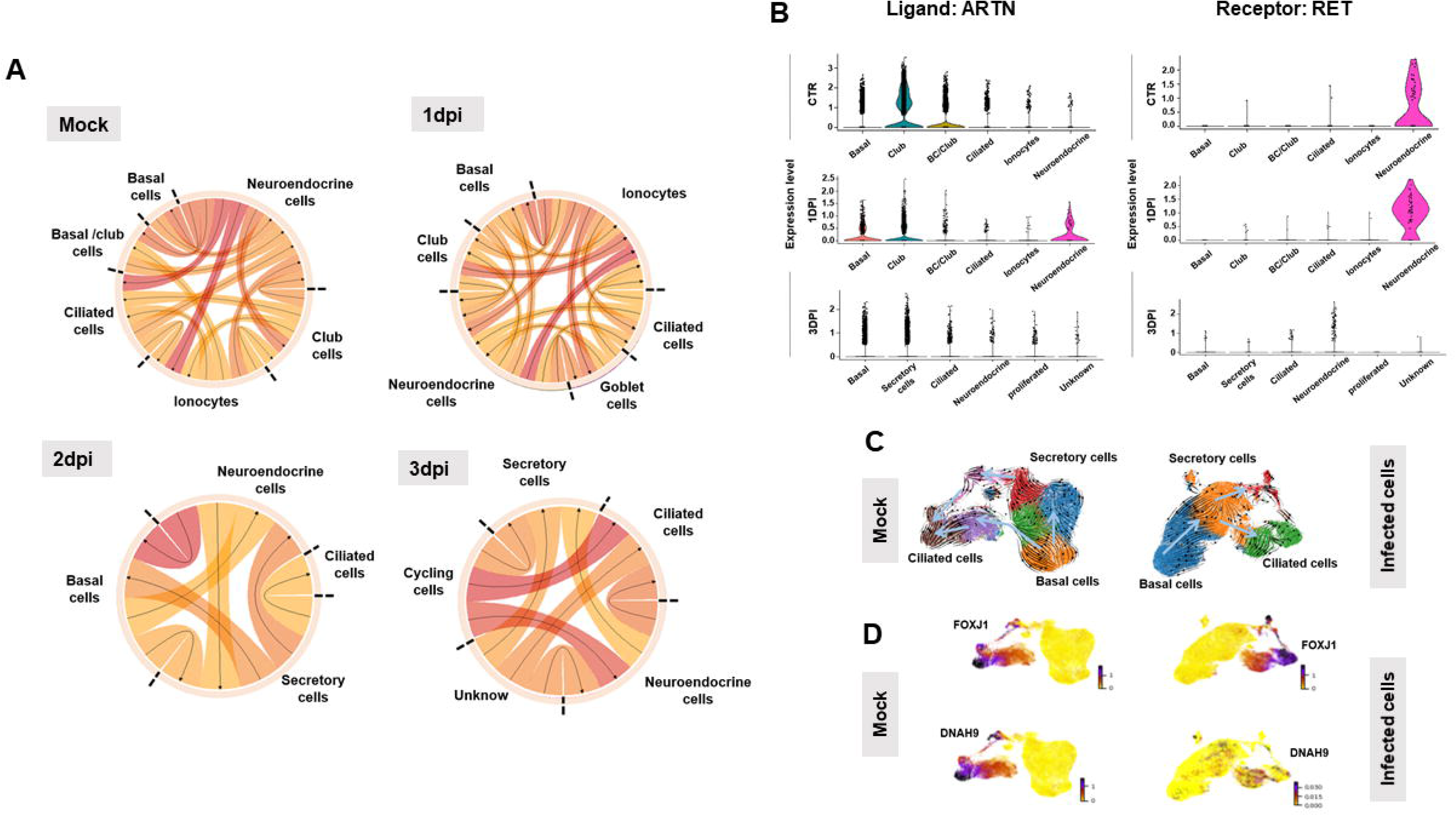
Paracrine interactions and single-cell trajectories in SARS-CoV-2-infected ALI cultures. (A) Circos plot showing potential interactions (ligands-receptors) made by various epithelial cell types in non-infected (Mock) and SARS-CoV-2-infected ALI cultures at 1, 2, and 3 dpi. The smallest number of paracrine interactions was observed in infected cultures at 2 and 3 dpi. Arrows point to the receptors. (B) Violin plots showing the expression of the ligand *ARTN* and its receptor *RET* in the various airway cell types identified by scRNA-seq before (CTR) and after SARS-CoV-2 infection (1 and 3 dpi). (C) The velocity field revealed two distinct tracks used by basal cells to form ciliated cells in non-infected ALI cultures (mock), but not in SARS-CoV-2-infected cultures (2 dpi). (D) Left, gene expression of known markers of premature (*FOXJ1*+) and mature ciliated cells (*DNAH9*+) is visualized on the t-SNE plot in the mock sample. Right, the t-SNE plot shows the absence of mature ciliated cells (*DNAH9*+) at 2 dpi.

### RNA velocity reveals discrepancies between non-infected and infected epithelium in the ALI model

To further characterize the ALI model transcriptional dynamics in response to virus infection, the single-cell RNA velocity was measured in SARS-CoV-2 infected and non-infected epithelial ALI cultures using a dynamic model of transcriptional state based on unspliced and spliced transcript counts [26]. Cell trajectory analysis revealed that non-infected ALI epithelial cells displayed two distinct bifurcation points through two different epithelial transition states. The first bifurcation point included basal cells (*TP63*-/*KRT5*+) that directly differentiate into ciliated cells, and the second bifurcation point mainly included basal cells (*TP63*+/*KRT5*+) that preferentially differentiate into secretory cells and then into ciliated cells. This placed secretory cells in an intermediate position between basal cells and mature ciliated cells (*DNAH9*+) (Figure 5C). Velocity ordering analysis in infected cells revealed a change in the ciliated cell differentiation trajectory. Indeed, basal cells were all oriented to differentiate into ciliated cells through a secretory cell state. Moreover, *DNAH9*+ infected cell mapping within ALI epithelial cells revealed that ciliated cells were more susceptible to infection compared with other cell types, like *DNAH9+* mature ciliated cells that are preferentially eliminated during SARS-CoV-2 infection (Figure 5D). Collectively, the dynamic ALI airway model provides a useful tool for *in vitro* studies on SARS-CoV-2/other virus infection and antiviral testing.

### Potential miRNA regulators of the epithelial cell intrinsic response to SARS-CoV-2 infection

The GenGo Metacore software was then used to identify miRNAs that regulate the epithelial cell response genes after SARS-CoV-2 infection in the ALI and iALI models (i.e. potential antiviral targets). This analysis identified 167 miRNAs that regulate the key epithelial cell intrinsic genes deregulated upon viral infection (Figures 3A-B). Among these miRNAs, 44 were regulators of interferon response genes and 123 regulated inflammatory response genes (complete list of miRNAs in Supplementary Table S6). More than 54% of the interferon gene targets were regulated by more than one miRNA. For instance, *SOCS3* and *MX2* were targeted by 24 and 5 miRNAs, respectively (Figure 6A). The inflammatory genes regulated by the highest number of miRNAs were *MMP9* (*n* = 29 miRNAs), *NFKBIA* (*n* = 19), *MMP13* (*n* = 19), *CSF1* (*n* = 12), *FOSL1* (*n* = 11) and *LOX* (*n* = 11) (Figure 6B). *MIR34a* was identified as a potential regulator of *MMP9*, *NFKBIA*, *FOSL1*, *CXCL10*, whereas *MIR203* and *MIR19* were common regulators of *NFKBIA, MMP13, SOCS3*. *MIR138-5p* and *MIR-326-3p* are validated regulators of *ISG15* and *ISG20*. *MIR650, MIR541-3p* and *MIR302d-3p* regulated several interferon□stimulated genes, including *MX1*, *MX2, IRF7* and *IRF9*. *CSF1* and *LOX* were targets of *MIR130a* and *MIR29*, respectively. These miRNAs can attenuate the inflammatory response and inhibit key coagulation cascade factors, thus preventing inflammatory epithelium damage. Therefore, they are potential miRNA-based therapeutics against COVID-19.

**Figure 6:**
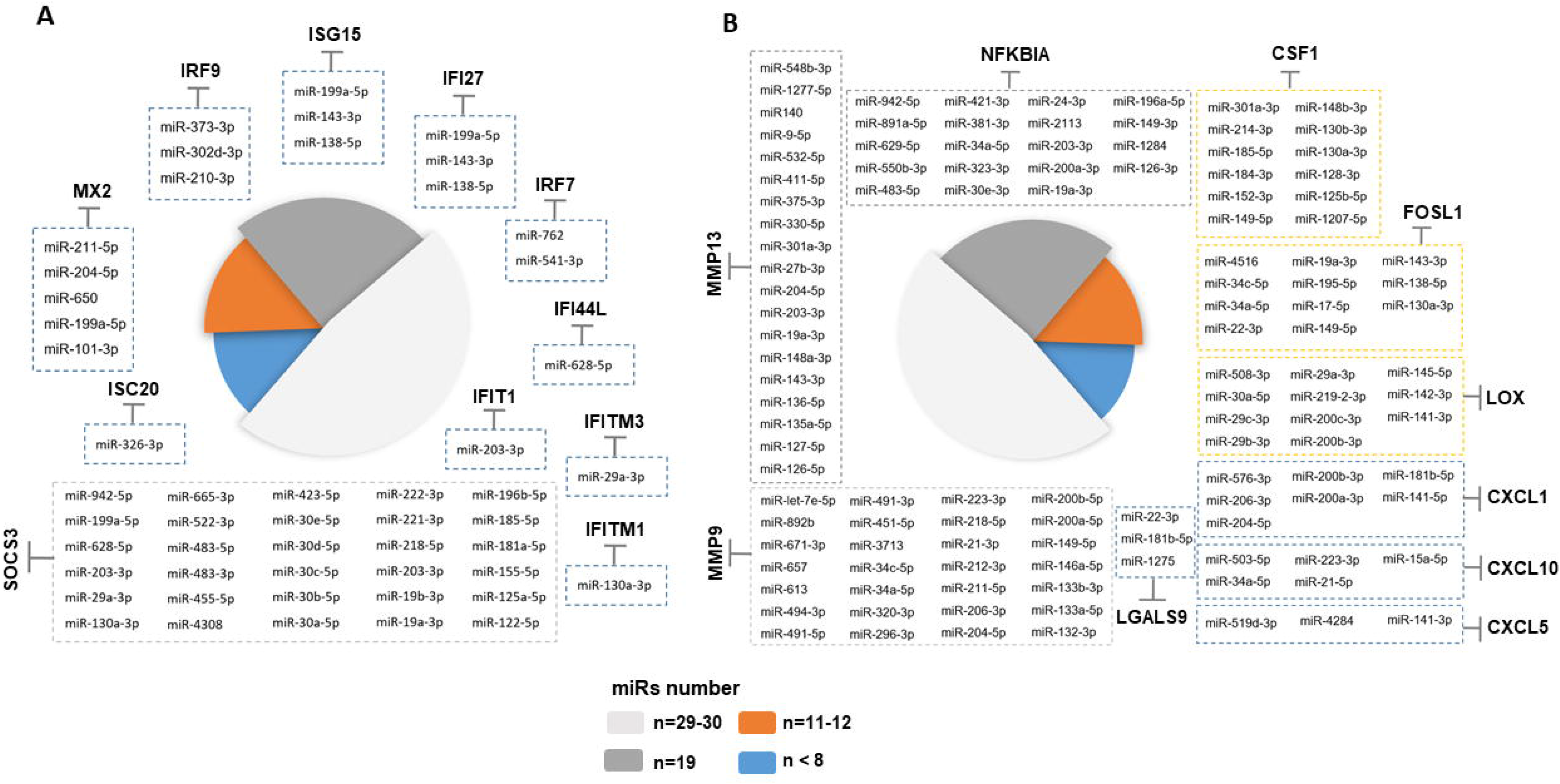
Identification of miRNA targets as potential COVID-19 therapeutics. Schematic representation of potential miRNAs that target some genes implicated in the interferon and inflammatory responses upon SARS-CoV-2 infection of ALI cultures. Only the miRNAs experimentally validated are represented. In the figure change into: Number of miRNAs targeting a gene.

## Discussion

The human airways are lined by an epithelium with three abundant specialized cell types (basal, secretory, and multi-ciliated cells) and some rare cell types (neuroendocrine cells, tuft cells and ionocytes) [35–37], on a stroma composed of mesenchymal cells, smooth muscle cells and immune cells [38]. Primary HBECs cultured in ALI conditions can be used to study airways *in vitro*. More recently, our group and others have generated functional airway epithelium from human iPSCs (iALI models) with a marked similarity to the airway epithelium *in vivo* [6,8,39–41]. As expected, scRNA-seq analysis of ALI and iALI models showed that immune cells, which are found in biopsy/brushing-derived primary cultures, were absent in these models. Conversely, the stromal component was exclusively found in ALI and iALI models, suggesting a role in the epithelium maintenance. Of note, neuroendocrine cells were mostly identified in the iALI model, compared with the ALI model. Our analysis characterized rare cell types, such as ionocytes, that strongly express *FOXI1* and *ASCL3* [36,42].

SARS-CoV-2 infection in ALI and iALI models induces virus-linked epithelial disruption, loss of mature ciliated cells, and triggers intrinsic immune responses. Despite the differences in terms of cell composition, the ALI and iALI models exhibit relevant proportions of airway cell types, express virus receptors (ACE2, CDHR3, CD55), and have been used to model various mechanisms of SARS-CoV-2 pathogenesis [43]. These models provide a suitable and reliable platform for researchers to study SARS-CoV-2 pathogenesis during infection.

In this study we analyzed the bulk and single-cell transcriptomes of ALI and iALI cultures infected with SARS-CoV-2. This highlighted the emergence of pro-inflammatory and interferon signatures in which epithelial cells activated the expression of several cytokines, chemokines, interferon, and downregulated ECM-related genes in response to SARS-CoV-2 at different times after infection. Most of these genes play essential roles in virus control and also in disease development. For instance, many chemokines (*CCL2*, *CCL3*, *CCL20*, *CXCL1*, *CXCL3*, *CXCL10*, *IL-8*) associated with inflammatory responses were expressed in the ALI and iALI models. *CCL2*, *CXCL10* (called *IP-10*) and *IL-8* are associated with airway inflammation, and high serum levels of these chemokines were found in patients with severe SARS [44]. *CCL3* also is involved in viral infections [45]. Our detailed analysis of the transcriptional response to SARS-CoV-2 infection showed that ALI and iALI cultures produced an unbalanced cytokine response, with preferential upregulation of genes encoding cytokines (for instance *IL-6*, *IL-1*β and *IL-33*) that are mainly implicated in the defense against extracellular aggression. Several studies showed that IL-6 serum levels are increased in patients with COVID-19 [46]. IL-6 is involved in acute inflammation due to its implication in controlling the acute phase response [47]. IL-6 production is increased by TNF-α and IL-1β [48]. In animal models of SARS-CoV-infection, TNF activity neutralization provides protection against SARS-CoV morbidity and mortality [49]. A large number of data showed the role of interferons in SARS-CoV2 infection. Interferons exercise their biological functions by regulating the expression of interferon-stimulated genes (ISGs). ISG upregulation has been described in various cells from patients with severe COVID-19 [50,51]. Here, we found that infection with SARS-CoV-2 induced a strong interferon response in ALI and iALI cultures, marked by high expression of *ISG15* (key factor in the innate immune response to SARS-CoV-2 infection), *ISG20* (with antiviral activity against RNA viruses), *IRF-7* (the master regulator of interferon responses) [52], and of several ISGs (*IFITM1, IFITM3*, *IRF9*, *IFI27*, *OAS2*, *MX1*, *MX2*, *SOCS3*) involved in the regulation of the host defense responses to the virus [53]. This suggests that *in vitro*, ALI/iALI epithelium can develop an intrinsic response to SARS-CoV-2 infection focused on the activation of the interferon pathways. This is in line with a study showing strong expression of many ISGs in the respiratory tract of patients with COVID-19, supporting the idea that interferon-mediated immune response plays a key role in SARS-CoV-2 infection control [51].

Although SARS-CoV-2 pathogenesis has not been fully understood, it seems that excessive immune responses play a key role. Evidence suggests that immune response deregulation causes lung damage [54]. Here we found that in infected ALI and iALI cultures, chemokines and cytokines, including IL-6, IL-1β, interferon and TNF, were upregulated to coordinate all aspects of the immunogenic response to SARS-CoV-2 infection. Additionally, SARS CoV2-related pneumonia with severe respiratory failure is characterized by enhanced ECM [55]. Similarly, in infected ALI cultures, cells expressing *MMP9*, an enzyme that participate in ECM remodeling, were markedly increased. The molecular pathways involved in *MMP-9* regulation during SARS-CoV2 infection are not known. Ueland et al. suggested that *MMP-9* may be an early indicator of respiratory failure in patients with COVID-19 [56], and Hsu et al. reported an important increase of MMP-9 concentration in the plasma of patients who developed acute respiratory distress syndrome [57]. Conversely, in infected ALI cultures, cells expressing ECM-related genes (e.g. *VIM*, *ITGB1*, *ITGB6*, *GJA1* and *PLOD2*) were decreased compared with control cultures. Vimentin and integrin are critical targets for SARS-CoV-2 host cell invasion [58,59]. Identifying the molecular mechanisms that lead to their regulation will be pivotal to understand their role in epithelium damage and reparation during SARS CoV2 infection.

Another important question we addressed concerned the miRNA role in the regulation of genes deregulated in infected ALI/iALI cultures. Recent investigations revealed that miRNAs are implicated in viral pathogenesis by altering the miRNA-modulated host gene regulation or the host immune system [60]. The variation of miRNA levels during virus infection and their role in modulating SARS-CoV-2 infection in human cells have been well described [17]. Thus, miRNA-antagonists or -mimics could be used to develop new therapeutic strategies for the treatment of patients with COVID-19 [61]. The anti-viral response by miRNAs may implicate the regulation of their mRNA targets that participate in the cellular response to viral infection. Interferon signaling was one of the primary effectors against SARS-CoV2 infections in both ALI and iALI models. Moreover, genes encoding members of this pathway are direct targets of many miRNAs. For instance, *MIR138* regulates *ISG15* expression by direct binding to its 3′ untranslated region (UTR) [62]. Similarly, *MIR326-3p* reduces the activity of an *ISG20* 3′ UTR luciferase reporter [63]. *MIR650*, *MIR29a-3p* and *MIR130a-3p* regulate other ISGs, such as *MX1*, *MX2*, *IFITM3* and *IFITM1* [64–66]. Moreover, many miRNAs target the 3’UTR site of matrix metalloprotease-encoding genes. For instance, *MIR34a* and *MIR19* target *NFKBIA* [67] (critical for SARS-CoV-2 entry in the cells) and *MMP9* [68] (macrophage-derived biomarker associated with inflammation), whereas *MIR130a* targets the 3’UTR of pro-inflammatory metabolite genes (e.g. *LOX* [69] that interacts with SARS-CoV-2) [70]. Altogether, different culture models are necessary to validate the biological effect of these candidate miRNAs to SARS-CoV-2 infection and to understand the interactions between host ISGs, cytokines and miRNAs. More investigations are needed to determine the expression profiles of key miRNAs during viral infection, to better predict disease severity, and to develop new therapeutic options based on these miRNAs for COVID-19 treatment and/or prevention.

## Conclusions

Here, we characterized of two ALI and iALI airway models to understand SARS-CoV-2 infection pathogenesis. We identified the inflammatory and interferon profiles induced in these models in response to SARS-CoV-2 infection. Understanding the long COVID19 effects remains a challenge and will require also these models.

## Funding

Supported by grants from the University Hospital of Montpellier (Appel d’offre interne, projet CILIPS 9174, projet INVECCO), the association Vaincre la Mucoviscidose (Grant #RIF20170502048), the Fondation pour la Recherche Médicale (Grant #FDM20170638083), the Labex Numev (ANR-10-LAB-20), and Astrazenca. The acquisition of the Cell-Discoverer 7 LSM900 microscope was funded by FEDER-FSE Région Occitanie throughout the MIP-FISH project. Part of this research project was supported by the CNRS INSB funding through the VIROCRIB program.

## Author contributions

Conceptualization and supervision: S.A., A.B and J.D.V.; Resources, A.B., J.D.V.; Data analysis and interpretation: S.A., A.B, A.N., S.W., N.G., D.M., A.B., J.D.V; Collection and/or assembly of data: S.A., E.A, F.F., C.B., A.P., I.V., D.M., A.B., J.D.V; Validation, L.M., N.G., D.M.; Writing – Original Draft, S.A, A.B, J.D.V.; Writing – Review & Editing, S.A., E.A, L.M., D.M., A.B., J.D.V.. Scientific and technical support: all authors listed have made a substantial, direct, intellectual contribution to the work, reviewed the final manuscript, and approved it for publication

## Ethics declarations

### Ethics approval and consent to participate

Subjects were recruited at Arnaud de Villeneuve hospital, Montpellier, France, under study protocols approved by the ethics committee - RRR study, NCT02354677, 2013-A01405-40 and INVECCO study, NCT03181204, 2017-A00252-51. All patients have been informed and agreed to participate by signing written consent forms.

### Conflict of interest

A.B. reports grants, personal fees, non-financial support and other from AstraZeneca; J.D.V. reports personal fees and other from Stem Genomics, personal fees and other from MedXCell Science, personal fees from Gilead, personal fees from Celgene, outside the submitted work. In addition, J.D.V. and S.A. have a pending patent EP20150306389. S.A. reports personal fees and other from Stem Genomics, outside the submitted work. E.A, L.M., A.N., F.F., C.B., S.W., A.P., I.V., D.M., declare no conflict of interest.

### Data set availability

Publicly available data were obtained from GEO datasets: GSE147507 and GSE166766. Interactive exploration tools: https://www.covid19cellatlas.org/index.healthy.html and https://cellxgene.cziscience.com/d/cellular_census_of_human_lungs_bronchi-17.cxg/. The gene expression profile from publicly RNA-seq data can be browsed with an interactive web-tool at: https://crem.shinyapps.io/iAEC2infection/ (GSE153277).

## Supplementary Tables

**Table S1:** Primer pairs used for validation by RT-qPCR

**Table S2:** List of the 515 genes specific to the SARS-CoV-2 bulk RNA-seq signature (transcriptome analysis by Blanco-Melo, D et al. [20]).

**Table S3:** List of the top GO categories identified by Ingenuity Pathway Analysis using the 515 genes listed in Table S2.

**Table S4:** List of the 146 enriched canonical pathways identified in the SARS-CoV-2 bulk RNA-seq signature.

**Table S5:** Lists of genes targeted by the upstream regulator’s *TNF*, *IL1A* and *F2* identified using IPA Upstream Regulator Analysis.

**Table S6:** Exhaustive lists of miRNAs that are putative regulators of genes implicated in interferon and inflammatory responses upon SARS-CoV-2 infection of ALI cultures retrieved by GenGo Metacore.

## Supplementary Figure

**Figure S1:** Mechanistic networks showing the interactions between upstream regulators (*TNF*, *IL1A*, *F2*) and their target genes identified in the SARS-CoV-2 bulk RNA-seq signature. The IPA upstream regulator analysis is based on the expected causal effects between upstream regulators and targets.

